# Quantitative RNA spatial profiling using single-molecule RNA FISH on plant tissue cryosections

**DOI:** 10.1101/2024.04.09.588031

**Authors:** Xue Zhang, Alejandro Fonseca, Konstantin Kutashev, Adrien Sicard, Susan Duncan, Stefanie Rosa

## Abstract

Single-molecule fluorescence *in situ* hybridization (smFISH) has emerged as a powerful tool to study gene expression dynamics with unparalleled precision and spatial resolution in a variety of biological systems. Recent advancements have expanded its application to encompass plant studies, yet a demand persists for a simple and robust smFISH method adapted to plant tissue sections. Here, we present an optimized smFISH protocol (cryo-smFISH) for visualizing and quantifying single mRNA molecules in plant tissue cryosections. This method exhibits remarkable sensitivity, capable of detecting low-expression transcripts, including long non-coding RNAs. Integrating a deep learning-based algorithm in our image analysis pipeline, our method enables us to assign RNA abundance precisely in nuclear and cytoplasmic compartments. Compatibility with Immunofluorescence also allows RNA and endogenous proteins to be visualized and quantified simultaneously. Finally, this study presents for the first time the use of smFISH for single-cell RNA sequencing (scRNA-seq) validation in plants. By extending the smFISH method to plant cryosections, an even broader community of plant scientists will be able to exploit the multiple potentials of quantitative transcript analysis at cellular and subcellular resolutions.

## INTRODUCTION

Gaining insights into the spatiotemporal changes in gene expression requires precise quantification and acquisition of high-resolution spatial distribution data for RNAs within cells and tissues. RNA i*n situ* hybridization (ISH) stands as a classical technique, enabling scientists to map RNA within intact cells or tissues with spatial precision. This is achieved through the use of probes containing nucleic acid sequences that complement a specific target RNA sequence (Pardue & Gall, 1969; Cox *et al*., 1984). ISH probes can be tagged with a variety of detectable molecules, such as radioisotopes (Harrison *et al*., 1973), biotin (Hutchison *et al*., 1982; Brigati *et al*., 1983), digoxigenin (Heiles *et al*., 1988; Panoskaltsis-Mortari & Bucy, 1995) and fluorescent dyes (Bauman *et al*., 1980; Singer & Ward, 1982). However, the traditional ISH techniques suffer from limited spatial resolution and are primarily qualitative in nature.

Single-molecule fluorescence *in situ* hybridization (smFISH) is a more recently developed variation of FISH that provides single-molecule resolution (Femino *et al*., 1998; Raj *et al*., 2008). It uses a set of 25-48 short DNA oligonucleotides (18-22 mers) to bind specific sequences in the target RNA, enabling the visualization and quantification of individual RNA molecules within intact cells and tissues at high spatial resolution. While smFISH has been successfully used in a variety of model organisms, including bacteria (Skinner *et al*., 2013), *C. elegans* (Raj *et al*., 2008), and mammalian cells (Lyubimova *et al*., 2013), its application within diverse plant tissues has been constrained by technical complexities arising from inherent autofluorescence and light-scattering components in cell walls and vacuole-rich tissue structures. The adaptation of smFISH to the plant model species *Arabidopsis thaliana* in 2016 marked a pivotal development, enabling the imaging of individual RNA molecules within fixed and squashed root meristem cells (Duncan *et al*., 2016, 2017). Since then, more smFISH and other newly developed ISH methods have gradually emerged for studying gene expression in plants (Yang *et al*., 2020; Solanki *et al*., 2020; Nobori *et al*., 2023; Huang *et al*., 2023). We have also recently optimized smFISH for whole-mount intact plant tissues (Zhao *et al*., 2023). Despite the success of whole-mount smFISH in several tissues, this method is not ideal for very thick specimens and physical sectioning may be required when dealing with plant tissues. The smFISH application to plant tissue sections has been reported in paraffin-embedded samples (Huang *et al*., 2020) but this workflow remains lengthy. Moreover, the complex embedding and prolonged processes introduced an increased risk for RNA degradation. As such, demand still exists for an expedited and streamlined smFISH method for tissue sections in plants.

In this study, we present an optimized smFISH protocol for the visualization and quantification of single RNA molecules in plant tissue cryosections. This method can enable quantitative analysis of weakly expressed transcripts, such as long non-coding RNAs, and provide precise information regarding transcript distribution across diverse cell types. Moreover, by incorporating DAPI staining and a deep learning-based cell-segmentation pipeline, we achieved segmentation of sub-cellular compartments, allowing us to distinguish and quantify nuclear and cytoplasmic transcripts. Additionally, we demonstrate the compatibility of smFISH with immunofluorescence (IF) in tissue sections, enabling simultaneous visualization and quantification of RNAs and endogenous proteins within individual cells. Significantly, this study demonstrates for the first time the application of smFISH for scRNA-seq validation in plants, investigating spatial gene expression patterns in relation to cell identity. In summary, our study illustrates how smFISH can be used for scRNAseq validation and for the simultaneous detection of endogenous RNAs and proteins.

## Results

### Single-molecule RNA detection and quantification on *Arabidopsis* and barley tissue cryosections

Plant tissue imaging poses specific challenges because of its inherent high autofluorescence. Although clearing techniques have greatly improved our ability to observe plant tissues in whole-mount settings (Zhao *et al*., 2023), it remains difficult to fully exploit these techniques in some plant species due to tissue thickness and high autofluorescence. In this context, the use of tissue sectioning emerges as a valuable alternative method. In contrast to optical sectioning, physical sectioning provides the advantage of imaging thicker specimens without sacrificing the signal-to-noise ratio. The first step in the development of our smFISH protocol on cryosections (hereafter referred to as cryo-smFISH) involved obtaining high-quality sections that preserve both the morphological integrity of plant tissues and RNA molecules. To accomplish this, we adapted previously published protocols (Anjam *et al*., 2016; Stapel *et al*., 2018) for use in *Arabidopsis thaliana* (*Arabidopsi*s) root cryosections (**Figure 1A**). This included a cryoprotection step conducted before cryosectioning to protect tissue morphology and a re-fixation step to ensure RNA preservation. These adjustments, combined with the use of 10*μ*m sections, enabled us to obtain well-preserved samples consisting of a single layer of cells. While *Arabidopsis* roots are not thick specimens, they are extremely fragile, posing significant challenges to tissue structure preservation. Therefore, optimizing a cryosection protocol tailored to Arabidopsis roots serves as a valuable initial step in the development of a robust method that can be applied to other types of tissues. To optimize and validate our cryo-smFISH protocol in *Arabidopsis* root cryosections, we focused on detecting transcripts from the housekeeping gene *Protein Phosphatase 2A* (*PP2A*) as previously reported (Duncan *et al*., 2016, 2017, 2023; Zhao *et al*., 2023). The resulting cryo-smFISH images from both longitudinal and cross cryosections revealed distinct bright punctate dots representing *PP2A* mRNA molecules, visible through both epifluorescence and confocal microscopy (**Figure S1**), with the confocal images offering the advantage of showing clearer cell outlines. *PP2A* mRNA molecules displayed a ubiquitous distribution across various root cell types in the root meristem region, consistent with earlier research findings(Duncan *et al*., 2016; Zhao *et al*., 2023). To enable precise allocation of transcripts to individual cells, we stained cryosections with the cell wall dye Renaissance 2200 (SR2200) (Musielak *et al*., 2016; Zhao *et al*., 2023). Combined with our integrated image analysis pipeline (Zhao *et al*., 2023), this enabled the precise and quantitative allocation of *PP2A mRNA* molecules to individual cells (**Figure 1B, C; Figure S3A,C**). Treatment with RNase confirmed that the observed signals corresponded to true mRNA molecules (**Figure S2**). We then assessed the applicability of our cryo-smFISH protocol in *Arabidopsis* leaves. Despite the challenges associated with autofluorescence in plant tissues, our protocol successfully detected *PP2A* RNA signals in young *Arabidopsis* leaves (**Figure S3B, D**). To further demonstrate the versatility of our approach, we extended our investigation to the monocot *Hordeum vulgare* (barley) by detecting the housekeeping gene *HvGAPDH* in cryosections of roots and leaves. Transcripts of this gene were clearly detected and quantifiable in both tissues (**Figure 1B, D; Figure S4A, B**). As before, treatment with RNase A confirmed the specificity of our RNA signals (**Figure S4C**). These results demonstrate that cryo-smFISH can successfully be used to detect and quantify specific RNA molecules with high resolution in cryosections obtained from both model and crop plant tissues.

**Figure 1.**
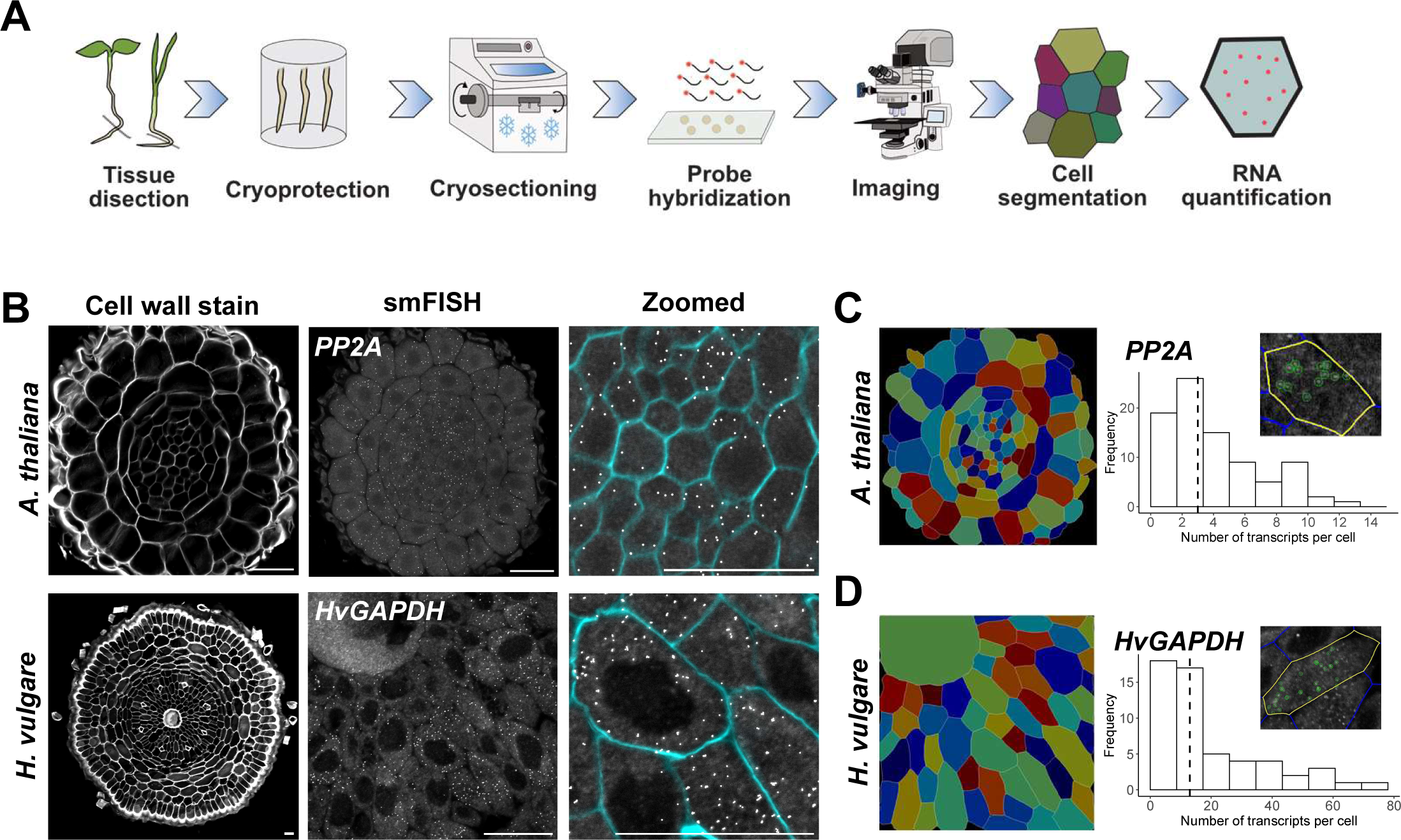
RNA detection and quantification on *Arabidopsis* and barley roots using cryo-smFISH. (A) Schematic illustration of cryo-smFISH detection and quantification workflow. (B) Single-plane confocal images of root cross-sections showing the detection of *PP2A* mRNA on *Arabidopsis* and *HvGAPDH* mRNA on barley. (left: cell contours visualized by cell wall staining with Renaissance 2200; middle: smFISH images with discrete white dots corresponding to individual RNA molecules; right: zoomed in image showing transcripts (white dots) and and cell walls (cyan)). Scale bars, 20μm. (C-D) Segmented cell masks for *Arabidopsis* (C) and barley cryosections (D) generated by Cellpose (left panels). Histograms display the quantification of cryo-smFISH signals for *PP2A* (C) and *HvGAPDH* (D) for the images depicted in B using FISHquant. The dashed line shows the median of the distribution. The insert images show examples of RNA detection in individual cells.

### Mapping the cell-type distribution of transcripts in *Arabidopsis* roots using cryo-smFISH

scRNA-seq has emerged as a powerful genomic approach for the detection and quantitative analysis of messenger RNA molecules at single-cell resolution. The pioneering collection of single-cell RNA sequencing (scRNA-seq) reports, applying high-throughput droplet-based technologies, has revolutionized our understanding of plant biology by revealing the transcriptional states of many different cell types (Denyer *et al*., 2019; Zhang *et al*., 2019; Wendrich *et al*., 2020; Shaw *et al*., 2021; Shahan *et al*., 2022; Otero *et al*., 2022). However, gene expression quantification results from scRNA-seq are not absolute but derived from computational algorithms. Hence, complementary methods are needed to validate its results. Furthermore, scRNA-seq outcomes lack spatial information as the tissue is disintegrated into individual cells before analysis (Shaw *et al*., 2021; Bawa *et al*., 2022). Complementary techniques, such as generating transgenic reporter lines and traditional ISH methodologies, are often utilized for scRNA-seq validation. Yet, these methods lack quantitative precision, fail to capture RNAs with particularly low expression, cannot visualize active transcription event or stress granules, and are unsuitable for determining kinetic transcriptional parameters. These limitations highlight the need for complementary techniques to validate scRNA-seq findings with cellular spatial information. Here, we demonstrate mapping the spatial distribution of transcripts in plant tissues using cryo-smFISH as a method suitable for scRNA-seq validation. As a proof-of-principle, we chose to study the nitrate transporter, *NRT1.9* (*AT1G18880*), which has been implicated in facilitating nitrate loading into the vascular subtype phloem cells in *Arabidopsis* roots (Wang & Tsay, 2011). To investigate cell-type specific expression of this gene, we re-analyzed scRNA-seq data from 10-day-old Arabidopsis root tips as previously described (Wendrich *et al*., 2020). Following clustering and cell type annotation step, we subset the dataset to specifically investigate cell types recognizable in tissue cryosections: epidermis, cortex, endodermis, pericycle, phloem, procambium, and xylem (**Figure 2A**). Analysis of differential gene expression (DGE) within these cell types indicated that *NRT1.9* is expressed preferentially the phloem, pericycle and procambium (**Figure 2A, B**). To confirm these results, we designed smFISH probes targeting exonic and intronic regions of *NRT1.9* RNA and performed cryo-smFISH on *Arabidopsis* root cross-sections (**Figure 2C-D**). The cryo smFISH results closely aligned with scRNA-seq expression pattern for *NRT1.9* (**Figure 2B, E, F**). A discrepancy was, however, noted in the procambium, where scRNA-seq indicated higher levels of expression. However, the trends in other cell types were very similar, and phloem cells consistently exhibited higher levels of expression in both scRNA-seq and cryo-smFISH (**Figure 2F**). Detailed inspection of the images revealed RNA foci of higher intensity (**Figure 2C**, indicated by orange arrows). Through DAPI staining, we confirmed that these brighter foci were predominantly present in the nucleus (**Figure S5**), potentially indicating active sites of transcription. Importantly, treatment with RNase A once again confirmed the specificity of our RNA signals (Figure **S6A-B**). Overall, these results highlight an important application of cryo-smFISH as a valuable method for precisely quantifying the transcription of genes of interest in a highly spatially resolved manner. Importantly, we demonstrated how cryo-smFISH can be used to validate and complement scRNA-seq findings at both qualitative and quantitative levels.

**Figure 2.**
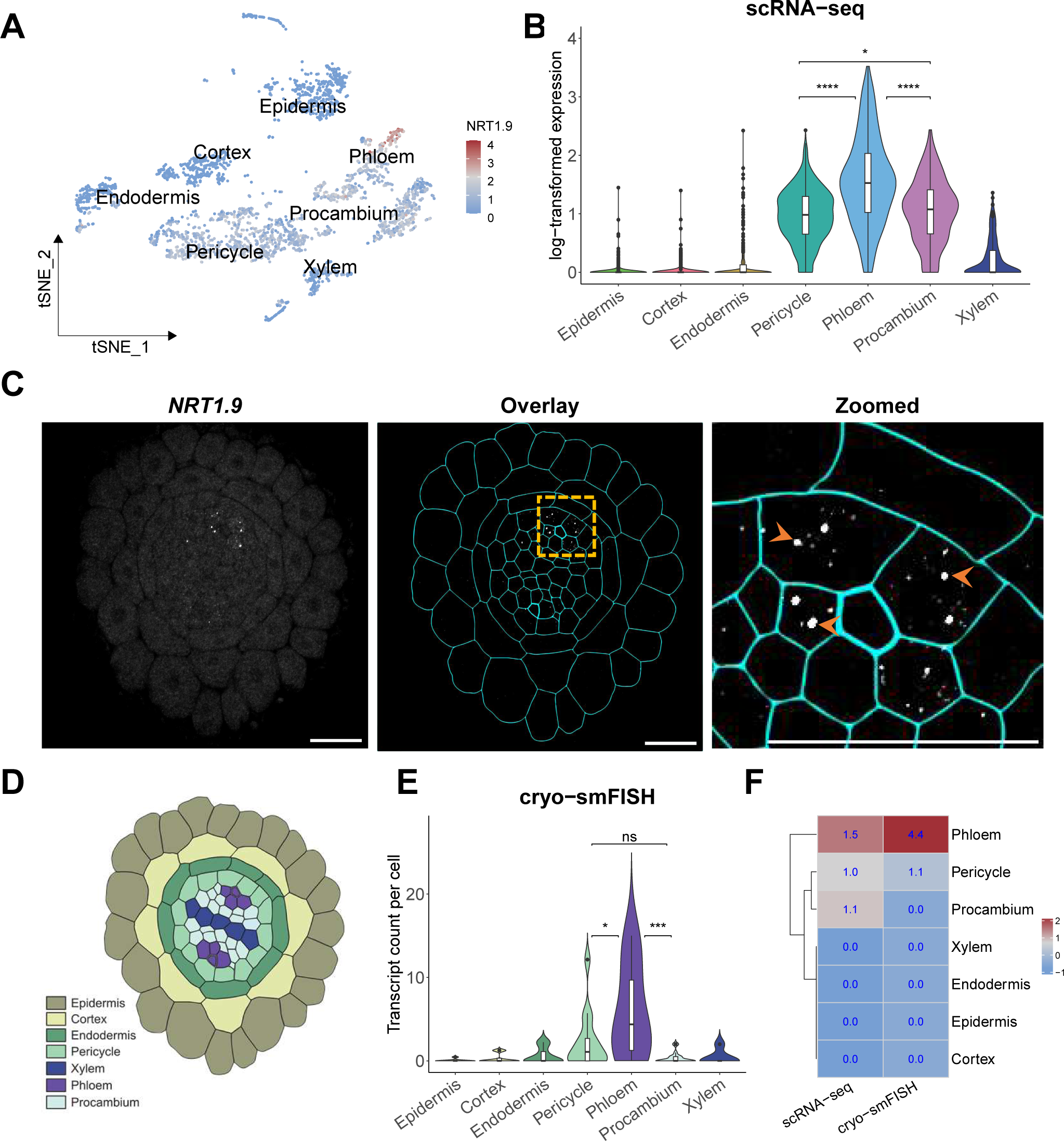
Mapping the cellular distribution of *NRT1.9* transcripts in Arabidopsis roots through scRNA-seq and cryo-smFISH. (A) The expression pattern of *NRT1.9* detected by scRNA-seq was visualized in t-SNE (t-distributed stochastic neighbour embedding). (B) ScRNA-seq analysis of *NRT1.9* expression in different cell types. (C) Representative images of cryo-smFISH showing *NRT1.9* transcripts on *Arabidopsis* root cross-section. Left panel: single-plane confocal image of cryo-smFISH for *NRT1.9;* Middle panel: cell outlines obtained from segmentation using cell wall dye SR 2200 (cyan) and *NRT1.9* RNA smFISH signals (white). Right panel: zoomed-in image showing *NRT1.9* transcripts (white dots) and cell walls (cyan). The orange arrows indicate bright *NRT1.9* RNA foci. Scale bars, 20μm. (D) Schematic of cell types in a cross-section of an *Arabidopsis* root depicted in C. (E) The violin plot illustrates quantification transcripts per cell from cryo-smFISH image (n=86 cells) depicted in C. Four replicates from same development stage show similar results. (F) Heatmap illustrates that both scRNA-seq and cryo-smFISH methods have identified *NRT1.9* as being highly expressed in phloem cells. The values displayed on the heatmap represent the median value of transcripts within each cell type, which have been normalized based on the cell quantity. Plotting area is scaled by width in violin plots B&E. The violin plots boxes present the interquartile range (IQR 25-75%), indicating the median values as a horizontal line. Whiskers show the ±1.58xIQR value. Statistical analyses were performed with one-way ANOVA followed by Tukey’s honestly significant difference (HSD) tests. A p-value greater than 0.05 indicates no statistical significance (ns), while p-values less than 0.05, 0.001, and 0.0001 were denoted by, *, and ***, respectively. In panel B and E, the comparison of p-values among pericycle, phloem, and procambium was displayed.

### Detection of long non-coding RNA distribution with subcellular resolution

Long noncoding RNAs (lncRNAs) are a class of RNA molecules with lengths over 200 nucleotides that have garnered increasing interest due to their potential regulatory roles in various biological processes (Lucero *et al*., 2020; Jha *et al*., 2020; Statello *et al*., 2021). In contrast to coding RNAs, lncRNAs are recognized for their pronounced tissue specificity and, often, relatively lower abundance (Djebali *et al*., 2012; Wang *et al*., 2014; Cabili *et al*., 2015; Rosa *et al*., 2016; Zhao *et al*., 2018). Despite predicted contributions to multiple various aspects of gene regulation, with subcellular localization being key to its function (Statello *et al*., 2021),the study of lncRNAs, especially at the single-cell level, remains limited (He *et al*., 2023). Here, we tested whether our cryo-smFISH method could be used to detect an uncharacterized lowly expressed lncRNA. *asSOFL1*, is the antisense lncRNA to *SOFL1*(Kim *et al*., 2022). We reanalysed bulk RNA-seq data from Arabidopsis wild type root tip(Choe *et al*., 2017). Among other genes, *asSOFL1* RNA was found to be weakly expressed but detectable within the root tip (**Figure S8B**). We next designed smFISH probes targeting *asSOFL1* transcripts and performed cryo-smFISH on *Arabidopsis* cross-sections. The results revealed the presence of bright dots, corresponding to *asSOFL1* transcripts (**Figure 3A**). Counting transcripts per cell revealed that, indeed, this lncRNA is weakly expressed with only a few transcripts detected per cell (**Figure 3B**). Despite its low expression, *asSOFL1* transcripts were found ubiquitously across various cell types within the root (**Figure 3C**). We then employed RNase A treatment, which confirmed that the bright spots observed corresponded to true RNA signals (**Figure S6C**). We additionally compared expression analysis of *asSOFL1* across root cell types obtained from scRNA-seq, bulk RNA-seq and cryo-smFISH data (**Figure S8**). This comparative analysis revealed that results obtained by cryo-smFISH are consistent with the other two methods, with similar trends also observed for *NRT1.9* and *PP2A*.

**Figure 3.**
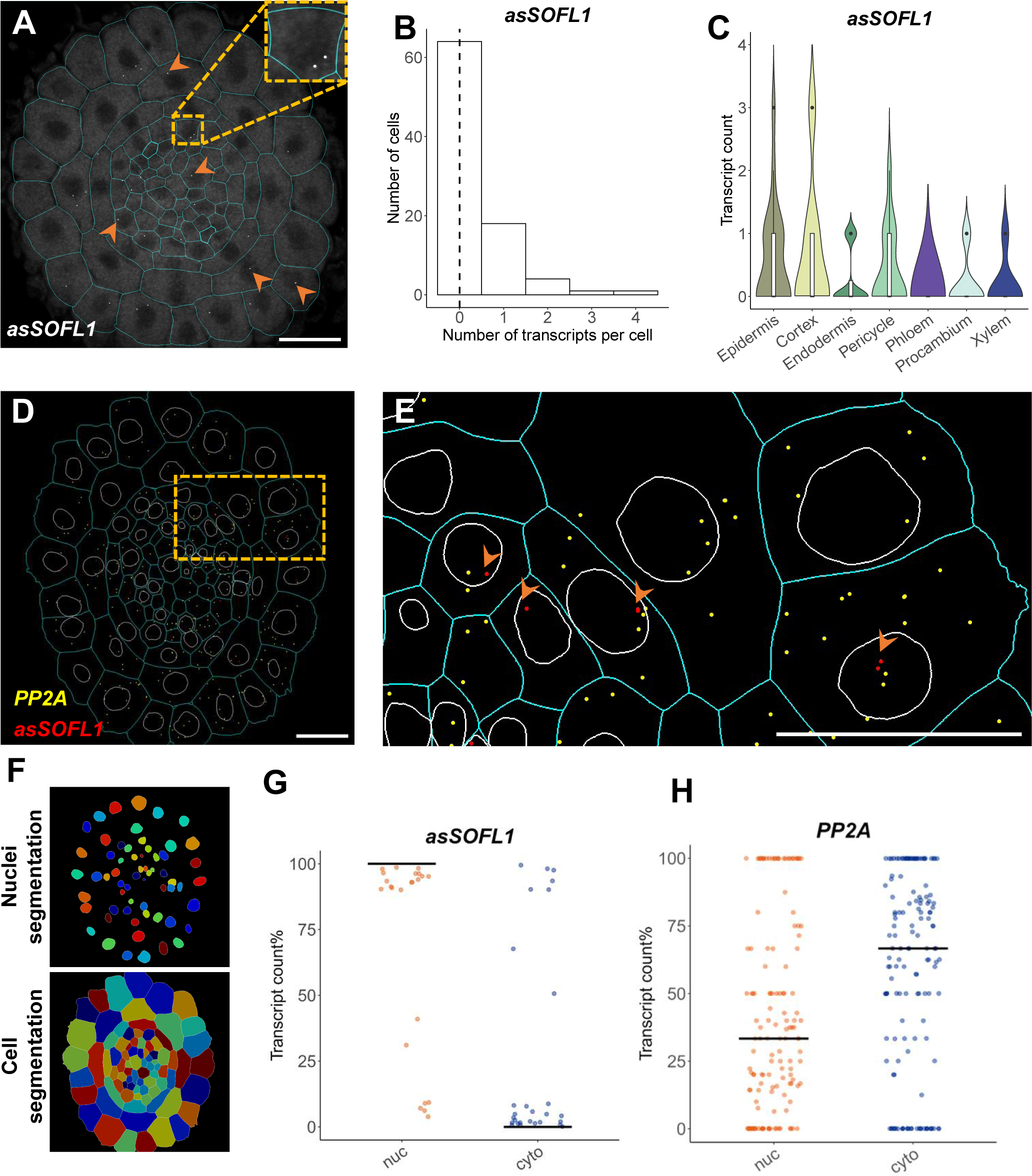
Cryo-smFISH enables the detection and quantification of long non-coding RNAs with subcellular resolution. (A) Representative cryo-smFISH image showing *asSOFL1* transcripts (white) on Arabidopsis root cross-section and cell outlines (cyan) obtained from segmentation using cell wall dye SR2200. Orange head arrows indicate *asSOFL1* RNA signals. (B) Histogram displays the distribution of *asSOFL1* transcript numbers in cells derived from image depicted in A. The dashed line indicates the median value of the transcript number detected within an individual cell. (C) Violin plot showing quantification of *asSOFL1* transcripts for different cell types, derived from image depicted in A. Experiments were repeated independently 5 times. Statistical analyses were performed with one-way ANOVA followed by Tukey’s honestly significant difference (HSD) tests. A p-value greater than 0.05 indicates no statistical significance (ns), while p-values less than 0.05, 0.001, and 0.0001 were denoted by *, **, and ***, respectively. Only significant differences were labelled. Boxes inside violin plots show the interquartile range (IQR 25-75%), indicating the median values as a horizontal line. Whiskers show the ±1.58xIQR value. (D) Detection of *asSOFL1* detection (red dots) and *PP2A* mRNA (yellow dots) using our image analysis pipeline. Cell and nucleus outlines obtained from segmentation using cell wall dye SR 2200 and DAPI respectively. (E) Zoomed-in image from the region highlighted in yellow in D. Orange head-arrows indicate *asSOFL1* RNA signals. (F) Cell and nuclei segmentation generated from DAPI channel image using Cellpose. (G, H) Quantification in percentage of transcripts in the nucleus (nuc) and cytoplasm (cyt) for *asSOFL1* (G, n=50 cells including nucleus for each group) and *PP2A* (H, n=201 cells for each group, 2 replicates are included) using jitter plots. The black horizontal line represent median value. Each dot indicates individual cell. Experiments were repeated independently 6 times. Scale bars, 20μm.

Subcellular localization patterns of lncRNAs can provide insight into their function (Cabili *et al*., 2015). To elucidate the subcellular localization of *asSOFL1* transcripts, we applied DAPI staining to our cryo-smFISH protocol. A visual inspection of images revealed that *asSOFL1* transcripts are mostly found in the nucleus (**Figure 3D, E**). In order to obtain a quantitative description in both nucleus and cytoplasm subcellular compartments, we modified the cell segmentation step in our image analysis pipeline. This was achieved by implementing Cellpose (Stringer *et al*., 2021), a deep learning-based algorithm for cellular segmentation (**Figure 3F**). Using two models trained specifically for segmenting cells and nuclei in *Arabidopsis* and barley tissue cryosections, we created cell masks and nuclei masks required as input files for subsequent transcript quantification in FISH-quant pipeline. With this adapted image analysis pipeline, we were able to automatically assign transcripts to either nucleus or cytoplasm compartments. Our cryo-smFISH results showed that almost 100% of *asSOFL1* transcripts are localized in the nucleus (**Figure 3G, Figure S7**). In contrast, over 70% of *PP2A* transcripts were found in the cytoplasm where translation occurs as expected for a coding gene (**Figure 3H**). Together, these results illustrate that our cryo-smFISH protocol can be used to detect lowly expressed transcripts, such as lncRNAs, and provide important data on their quantitative expression at the cellular and subcellular levels.

### Simultaneous RNA and Protein Detection by sequential smFISH-IF protocol

Exploring the subcellular localization of proteins in conjunction with gene expression quantification and mRNA localization can shed light on the complex interplay between transcriptional and post-translational gene regulation. To assess the compatibility of our cryo-smFISH technique with immunofluorescence (IF), we used *Arabidopsis* root cryosections and adapted a sequential smFISH-IF protocol as previously described (Rosa *et al*., 2016). Using probes against the *PP2A* mRNA, we first performed cryo-smFISH, followed by the IF protocol using an antibody against the histone marker H4Ac. The results revealed that the antibody staining worked well after cryo-smFISH protocol with uniform signals across all cells and good tissue preservation, allowing an easy overlay of cryo-smFISH and IF images (**Figure 4A**). This dual approach allowed us to not only visualize the presence of *PP2A* mRNA and H4Ac in the same cells but also quantiify their levels per cell (**Figure 4B, C**) and assess their abundance within the same cells and across different cell types (**Figure 4D**). Importantly, this sequential cryo-smFISH/IF protocol can be extended to crop species such as barley (**Figure 4B**). The development of this protocol will be of important application for colocalization studies especially for species for which transgenesis has not been achieved.

**Figure 4.**
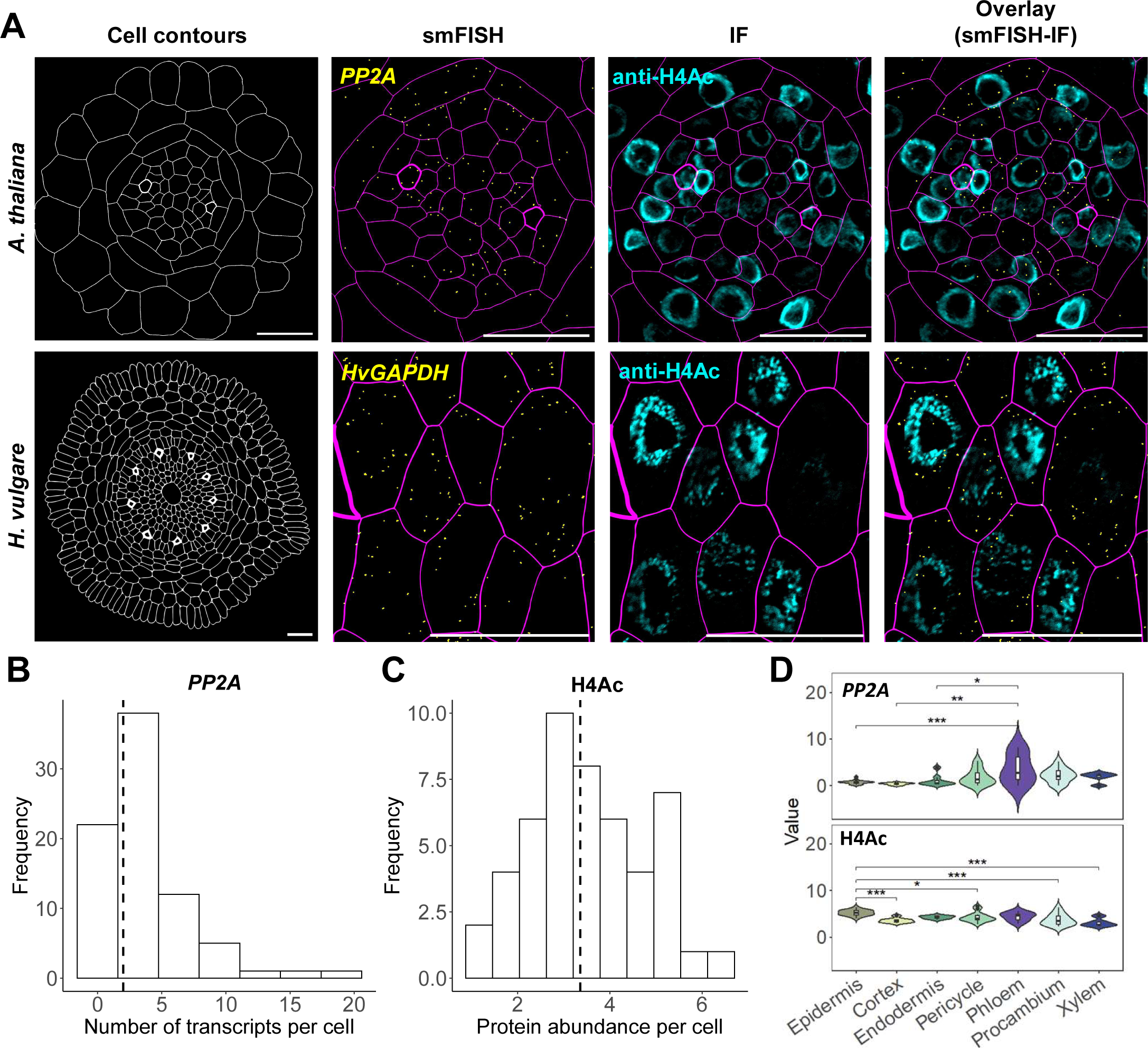
Sequential cryo-smFISH-IF protocol enables simultaneous detection and quantification of RNAs and endogenous proteins in single cells. (A) Images obtained using sequential cryo-smFISH-IF protocol on cross-sections of *Arabidopsis* and barley. (Cell wall outlines (white our magenta) obtained from segmentation using cell wall dye SR2200; Detected *PP2A* and *HvGAPDH* transcript (yellow); Detection of acetylayed histone H4, H4Ac (cyan)). Scale bars,20μm. (B, C) Histograms display the number of *PP2A* transcripts per cell (B) and the abundance of H4Ac endogenous protein levels per cell (C) in Arabidopsis root cross-sections. The dashed lines indicate median values. (D) Violin plot shows the distribution of *PP2A* mRNA and H4Ac protein levels within cell types from same Arabidopsis root cross-section. Values represent transcript number/protein intensity normalized by cell size. Boxes inside violin plots show the interquartile range (IQR 25-75%), indicating the median values as a horizontal line. Whiskers show the ±1.58xIQR value. Statistical analyses were performed with one-way ANOVA followed by Tukey’s honestly significant difference (HSD) tests. A p-value greater than 0.05 indicates no statistical significance (ns), while p-values less than 0.05, 0.001, and 0.0001 were denoted by *, **, and ***, respectively. Only significant differences were labelled. Experiments were repeated independently 4 times.

## Discussion

In this study, we introduce an optimized smFISH protocol designed specifically for plant tissue cryosections. Our cryo-smFISH method provides an effective approach to obtain the precise quantitative detection of RNA spatial distribution with single-cell and single-molecule resolution in plant tissue sections. This approach is relatively easier and faster compared to other sectioning methods such as the ones involving paraffin embedding. While cryosectioning poses greater challenges for tissue preservation compared to paraffin sections, it excels in maintaining the stability of biomolecules, including protein epitopes and nucleotides (Hira *et al*., 2019). Compared to our recently established smFISH for whole mount tissue, the choice of method will depend on the type of material and the final objective of the study. While whole-mount smFISH performs well in preserving the tissue structure it may yield inferior signal-to-noise ratios for thicker specimens. Furthermore, if the goal is to combine RNA with protein detection through immunofluorescence, cryosectioning offers a distinct advantage, as antibody penetration is notably more efficient in tissue cryosections. The use of thin sections also allows for the use of widefield microscopes, which can be invaluable when confocal microscopes are not readily available. Moreover, the use of widefield systems for cryo-smFISH samples can significantly expedite image acquisition compared to confocal imaging, while still allowing for excellent segmentation results.

The complexity of vascular tissue structure and development have historically made high-resolution gene expression profiling challenging. Detailed gene expression profiling has predominantly been explored through scRNA-seq (Wendrich *et al*., 2020; Chen *et al*., 2021; Otero *et al*., 2022; Kułak *et al*., 2023). Here, we employed cryo-smFISH to physically map and quantify the vascular tissue-specific gene *NRT1.9* within individual cells in a transgene-free manner. Our analysis, combining scRNA-seq with cryo-smFISH, highlights the significant application of smFISH as a powerful method for validating scRNA-seq data while complementing it with cellular or subcellular spatial information. This further reinforces the importance of expanding the use of smFISH for several plant tissues and preparations, particularly as the demand for scRNA-seq validation methods continues to rise.

Lastly, cryo-smFISH exhibits sensitivity in detecting lowly expressed genes, such as long noncoding RNAs, thereby expanding our capabilities to study RNA function at both cellular and subcellular levels. However, it’s worth noting some limitations. While some tissues, like barley roots, yield good cryosections relatively easily, others, such as leaves or *Arabidopsis* roots, may present challenges in maintaining tissue integrity and require careful optimization and practice. Additionally, like most smFISH methods, our protocol has limitations in detecting RNAs with lengths shorter than 600 nucleotides and co-detecting multiple RNA species (see **Table S1** for additional strengths and limitations of cryo-smFISH).

In summary, our cryo-smFISH method represents a valuable tool for conducting precise quantitative studies of single-molecule RNA within plant tissue sections in a highly spatial resolved manner. This innovative approach enables plant biology researchers to explore the complexity of transcriptional and translational products at both cellular and subcellular levels, thereby greatly expanding the scope of research possibilities in this field.

## Acknowledgements

We thank A. Menkis for initial cryostat technical support and Alexandre Berr for giving scientific feedback, and members of the Sicard and Rosa groups for discussion and comments on the article. This work was supported by Knut and Alice Wallenberg Foundation (KAW 2019-0062) and Carl Tryggers Stiftelse (CTS 18-325).

## Contributions

X.Z., A.S. and S.R. designed the research. X.Z. performed cryo-smFISH experiments; K.K. performed IF experiments; X.Z. analyzed scRNA-seq and bulk RNA-seq; A.F. and X.Z. performed image analysis; X.Z. and A.F. prepared figures; S.D. contributed to method optimization steps; X.Z. and S.R. wrote the manuscript. All authors commented and approved the final manuscript.

## Data availability

These supplementary files, and all data underlying the graphs and heatmaps presented can be accessed at https://github.com/xuezhang911/zhang_et_al_smFISH_cyrosections.

## Code availability

All custom code (R/Python scripts) can be accessed at https://github.com/xuezhang911/zhang_et_al_smFISH_cyrosections.

## Methods

### Plant materials and growth condition

Arabidopsis (*Arabidopsis thaliana*) Col-0 and Barley (*Hordeum vulgare*) were used in this study. Barley (*Hordeum vulgare*) seeds were a gift from Silvana Moreno (Department of Plant Biology, SLU – Uppsala, Sweden). Seeds were submerged with 1ml 70% ethanol in 2ml Eppendorf tubes and placed on the rotating wheel for 5-10mins, followed by three times wash with 95% ethanol. Then, ethanol was removed, and seeds were placed in the clean hood to dry.

After 2-3 days of stratification seeds were sown on Murashige and Skoog (MS) agar plates, the plates were transferred to a growth chamber with the following conditions: photoperiod of 16 hours day and 8 hours night and a temperature cycle of 22°C during the day and 20°C during the night. Arabidopsis root tips were collected 7 days after germination. Barley roots were collected 4 days after germination.

### smFISH probe design

Probes targeting the genes of interest were designed using online website from Biosearch Technologies: https://www.biosearchtech.com/stellaris-designer. The sequences of the probe were then subjected to quality control process using automated local blast R-script, available in github at: https://github.com/xuezhang911/zhang_et_al_smFISH_cyrosections/tree/main/smFISHprobes. The smFISH probes used in this study and respective fluorophores are shown in Table S2. The probes were diluted in TE buffer at a final stock concentration of 25 μM.

### Cryo-smFISH

#### I. Sample preparation

Tissues were dissected using a razor blade and fixed with 4% (m/v) paraformaldehyde (PFA) solution in nuclease-free 1xPBS buffer (pH8.0, Sigma-Aldrich, Cat.# AM9624). Following fixation, the tissues were subjected to a cryoprotection process with 34% sucrose in 1xPBS, followed by an overnight incubation with mild shaking in a mixture of equal parts 34% sucrose in 1xPBS and optimal cutting temperature compound (OCT; Leica Biosystems Cat.#14020108926) liquid at 4°C. Subsequently, the samples were exposed to pre-chilled OCT liquid for 30 minutes to an hour with mild shaking before preparing tissue blocks. Then, using tweezers, the tissues (either roots or leaves) were gently placed into 3D-printed Cryomolds (1cm x 1cm x 1cm), filled with pre-chilled OCT. These OCT-embedded tissue blocks were frozen, either by contact with dry ice or indirect contact with liquid nitrogen. Cryosection blocks were then wrapped in foil and stored at −80°C until cryosectioning. 10 μm (Arabidopsis and barley roots) or 5 μm (barley leaves) of cryosections cut by cryostat (Leica CM1850) were attached to selected polysine adhesion slides (Polysine^TM^ adhesion microscope slides Cat.# 48382-117) followed by air-dry for up to 20 mins and post-fixation for 10mins with 4% PFA at RT. Subsequently, after rinsing 3 times with 1xPBS, the slides with sections were subjected to permeabilization using methanol, ethanol, and clearsee solution: Xylitol powder10% (w/v; Sigma-Aldrich, Cat.# X3375), Sodium deoxycholate 15% (w/v; Sigma-Aldrich, Cat.# D6750) and Urea 25% (w/v; Sigma-Aldrich, Cat.# U5378) dissolved in RNase-Free Distilled Water (Qiagen, Cat.#10-977-015. The slides were then stored at 4°C overnight, ready for subsequent smFISH procedure.

#### II. Hybridization

The procedure is conducted as described previously with slight modifications (Duncan et al., 2016; Zhao et al., 2023). Briefly, slides prepared with cryosectioned samples were first rinsed 2-3 times with wash buffer (10% (v/v) nuclease-free; 20× SSC (Thermo Scientific, Cat. #AM9763); 10% (v/v) deionized formamide (Merck, Cat.#S4117); 0.1% (v/v) Triton X-100 (Sigma-Aldrich, Cat.#T8787). 100 μl of hybridization solution (10%(w/v) dextran sulfate; 10%(v/v) deionized formamide) with probes at a final concentration of 250 nM was added to each slide. Plastic coverslips were laid over the samples to prevent buffer evaporation and the probes were left to hybridize in a humid chamber at 37 °C overnight in the dark.

#### III. Post-hybridization and mounting

Plastic coverslips were gently removed and hybridization solution containing unbound probes was rinsed out with 2-time wash using wash buffer. Subsequently, slides were immersed in coplin jars containing wash buffer for up to 1 hour at 37 °C. DAPI (1:10000) or SR2200 dye (1:50000; Renaissance Chemicals Ltd., see Recipes Musielak et al. 2015;) in wash buffer was applied for 10-15 mins and 5-10 mins respectively. After 5-10 mins of washing with 2xSSC buffer, samples were mounted in freshly prepared GLOX mounting medium (0.4 % glucose; 10nM Tris–HCl, pH 8.0 (Invitrogen, Cat.#AM9856) and 2× SCC; 1/100(v/v) glucose oxidase (Sigma,Cat.# G2133-10KU) and 2/100 (v/v) mildy vortexed catalase catalase enzyme from bovine liver(Sigma, Cat.# C3155-50MG)) and sealed with nail polish.

### smFISH followed by immuno-fluorescence (IF)

We conducted cryo-smFISH and IF sequentially, cryo-smFISH protocol for RNA detection was performed first according to the earlier description, followed by immunofluorescence protocol after cryo-smFISH image acquisition. After imaging, the coverslips were gently removed and samples were then rinsed three times with 1xPBS solution. Samples were then incubated with the enzyme mix (2% Driselase, 1% Cellulase, 2% Macerozyme in 1x PBS) for 15 min in a humid chamber at 37 °C. After washing three times with 1xPBS, samples were incubated with 5% BSA blocking buffer in a humid chamber for 30 min at 37 °C. Subsequently, the samples were incubated with the H4ac antibody (Bio-Rad, AHP418) diluted 100 times in 5% BSA in a humid chamber overnight at 4 °C. The samples were then washed with PBST (0.1% Tween in 1xPBS) buffer by incubating in Coplin jars at 37 °C for 30 min. Then samples were incubated with secondary antibody (Agrisera, AS09633) diluted 200 times in 5% BSA in a humid chamber at 37 °C for 2 hours. The excess antibody was removed, and samples were washed with PBST in coplin jars at 37 °C in the dark for 30 min. Samples were then rinsed with 1xPBS and mounted in a drop of Vectashield (Bionordika, Cat.# VEH-1000) medium.

### Image acquisition

A Zeiss LSM800 inverted confocal microscope (Zen Black Software) was used for imaging through a 63X water-immersion objective (1.20 NA). We acquired z-stacks with 0.2 µm spacing. The following channel settings were used: DAPI/SR2200: 353nm excitation, 420– 470 nm emission; Quasar570: 548 nm excitation, 570–640 nm emission; Quasar670: 650 nm excitation, 665–715 nm emission; H4ac: 488 nm excitation, 500–550 nm emission.

For widefield microscopy, we acquired varying from 10 to 40 z-stack images with a cooled quad-port CCD (charge-coupled device) ZEISS Axiocam 503 mono camera through a 63X water-immersion objective (1.20 NA). The following channel settings were used: Quasar570, 533-558nm excitation, 561nm emission; Quasar670: 625-655nm excitation, 673nm emission; DAPI/SR2200: 335-383nm excitation, 465nm emission.

### Image analysis

To quantify the number of transcripts and protein levels per cell in an unbiased manner, we adapted the automated computational workflow utilized by Zhao et al., 2023. Our workflow mixes functions of four freely available software programs: Fiji (Schindelin *et al*., 2012), Cellpose (Stringer *et al*., 2021), FISH-quant (Mueller *et al*., 2013; Imbert *et al*., 2022), CellProfiler (Carpenter *et al*., 2006; Stirling *et al*., 2021) and python script. To conduct the analyses, we worked with separated TIF images for each channel: SR2200, DAPI, smFISH, or IF. Shifts and misalignments between channels and images acquired for the same section were corrected using the BigWarp tool in Fiji (Bogovic *et al*., 2016). The maximum z-projection or single plane confocal or wide field images were used for analysis.

### Cell and nuclear segmentation

Image segmentation was performed using Cellpose-trained algorithms (Stringer *et al*., 2021). For cell segmentation, using SR220 channel images as input, the ‘cyto’ algorithm was selected as pre-trained model for annotating individual cells within the tissue. To achieve accurate cell segmentations, a semi-supervised training for SR2200 channel images was implemented. Following annotation step, manual corrections and consecutive training cycles were applied until cell borders were precisely outlined. New models specifically segmenting cells of Arabidopsis or Barley tissue sections were generated, which were utilized to generate cell masks for subsequent transcription quantification step.

For the nucleocytoplasmic level analyses, we first trained a new segmentation model from the pre-loaded ‘nuclei’ algorithm or from cell segmentation model developed earlier from SR2200 images, using semi-supervised training as outlined above. As result, we established two models specifically segmenting cells with well-defined cell borders and nucleus within Arabidopsis or Barley tissue sections. These two models were thereafter utilized to create cell and nuclei masks for subsequent transcription quantification step.

### Image filtering and dot quantification

We employed MATLAB version of FISH-quant v3 for quantifying RNA dots in cryo-smFISH images (Mueller *et al*., 2013; Imbert *et al*., 2022). The software and its accompanying manual can be accessed on Bitbucket, provided by Florian Mueller: https://bitbucket.org/muellerflorian/fish_quant/src/master/.

The “Cell segmentation” tool was first utilized to generate text files containing FISH-quant recognized nuclei or cell outline coordinates from the Cellpose-generated masks. After the outline text file for a given image was imported, the loaded cryo-smFISH image was filtered by applying a Dual-Gaussian filter. For wide field cryo-smFISH image in figure S*7*, before being imported to FISH-quant, it was preprocessed with deconvolution lab2. The filtered image was then smoothed with a Gaussian Kernel. The Gaussian Kernel ratio required adjustment for each gene and in various tissue contexts.

Subsequently, we conducted pre-detection of dot in one selected filtered image. It was performed in the filtered image by defining threshold intensity and quality scores that fit precise fluorescent foci detection. Following this, detected foci were fitted using Gaussian fluorescence fitting based on the point-spread function (PSF). The settings were next implemented to run the analysis in batch mode.

False positives were removed by thresholding the Sigma-XY, amplitude, and pixel-intensity parameters, following the developer’s advice. Results were exported as tabulated files, indicating the number of transcripts per cell and per nuclei for every cell identifier.

### Image visualization

Cell or nuclei visualization: Cell and nuclei masks were created using Cellpose and imported into CellProfiler to generate cell or nuclei outlines that trace the contours of individual cells and their nuclei. These cell outline images were then imported into Adobe Illustrator. The high-resolution images with fine preservation of detailed tissue structures were finalized through image vectorization and a manual adjustment of the membrane thickness for phloem seize element cells from Arabidopsis and Barley root.

Cryo-smFISH dot visualization: an image displaying cryo-smFISH dots was generated either from the filtered image during transcript quantification step or from manually plotted dots based on the coordinates provided by the final quantification output. Then this image was overlaid with original cryo-smFISH RNA channel to enhance the visualization.

### Protein quantification

The immunofluorescence images were analyzed using Cellprofiler (Carpenter *et al*., 2006; Stirling *et al*., 2021). Cellpose-generated masks were imported and used to identify each nucleus or cell as an independent object. From the IF images, we computed both the total intensity and the mean intensity for each object, allowing us to determine protein levels either on a per-nucleus or per-cell basis. Results were exported as tabulated files.

### RNA or protein assignment to individual cells within cell type

To assign gene transcript numbers to specific cell types, we first obtained RNA mask and cell/nuclei mask. For RNA mask, we visualized RNA molecules by displaying their XY coordinates on a blank image. These coordinates representing the center of dots were obtained during transcription detection step. For cell/nuclei mask, we used Cellpose generated masks. Subsequently, we imported both RNA mask and cell/nuclei mask into CellProfiler, and the number of RNA molecules per individual cells was counted and displayed on cell/nuclei masks. Similarly, after we measure the protein intensity with CellProfiler, the intensity within individual cells was counted and displayed on cell masks. Consequently, images with well-defined tissue structures, cell/nuclei borders and RNA masks as graphical representations of the single-cell results were produced.

### scRNA-seq analysis

Normalized counts file in h5 format was downloaded from the Gene Expression Omnibus (GSE141730_aggregated_filtered_gene_bc_matrices.h5). Feature selection, dimension reduction, and clustering were performed as the original article described (Wendrich *et al*., 2020). Cell types were manually annotated using marker genes provided by the article. Then for data analysis in this article, we subset the whole Seurat dataset to only focus on the cell types including Epidermis, Cortex, Endodermis, Pericycle, Phloem, Procambium and Xylem cells.

### Bulk RNA-seq analysis

The fastq file for WT was downloaded from the NCBI study GSE96945. After rRNA removal by SortMeRNA, adaptors were trimmed by Trimmomatic. The gene raw counts matrix was obtained with pseudo-align software Kallisto. RNA-seq reads were normalized by transcript per million.

