## Supplemental Figures for "Quantitative RNA spatial profiling using single-molecule RNA FISH on plant tissue cryosections"

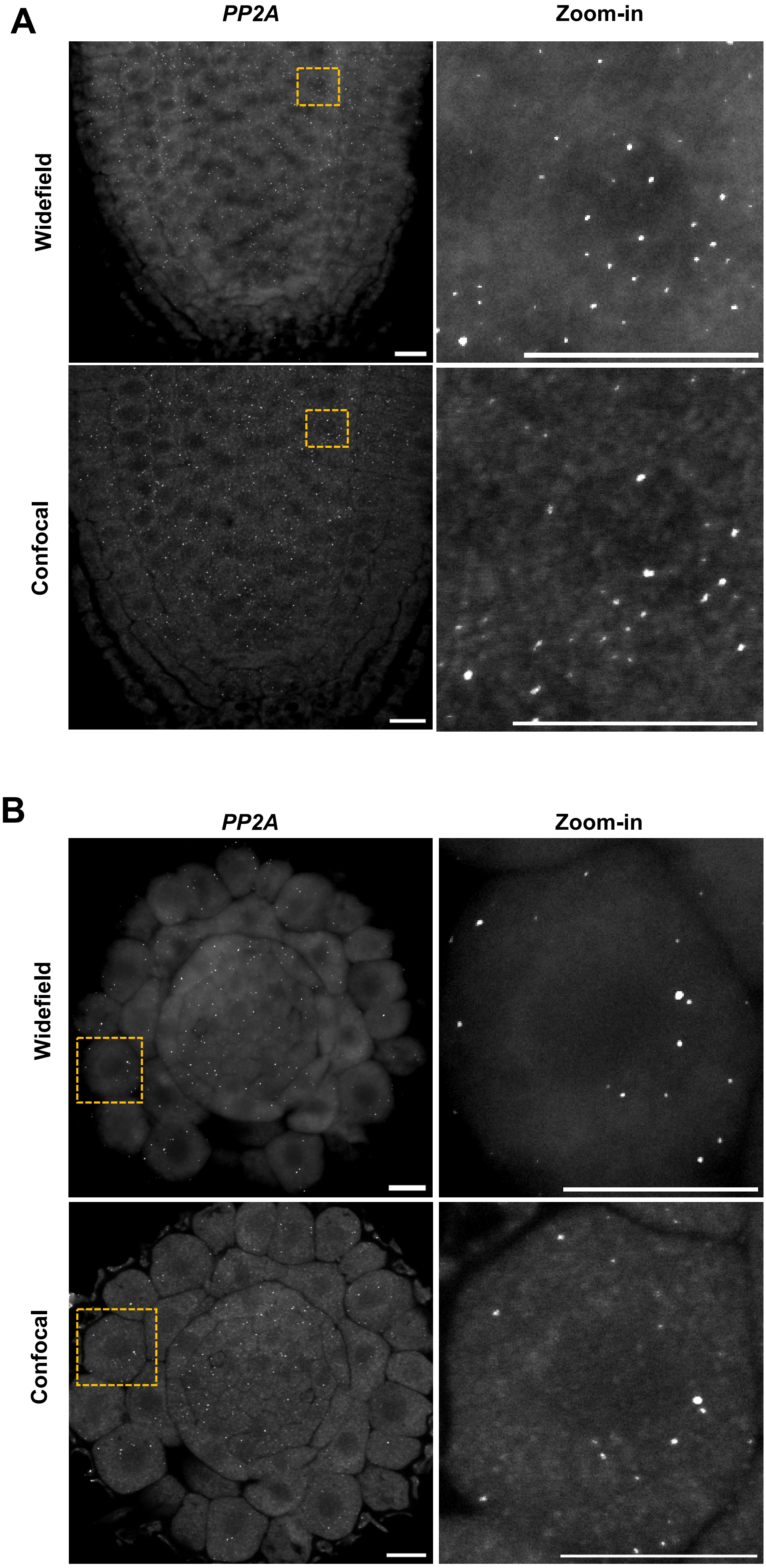


**Figure S1. smFISH on *Arabidopsis* root longitudinal and cross-cryosections.**
Detection of *PP2A* mRNA using smFISH on *Arabidopsis* root longitudinal (A) and cross-sections (B). Zoom-in images from regions highlighted in yellow on the left panel. Discrete white dots above the diffuse background correspond to individual RNA molecules. Scale bars, 10µm.

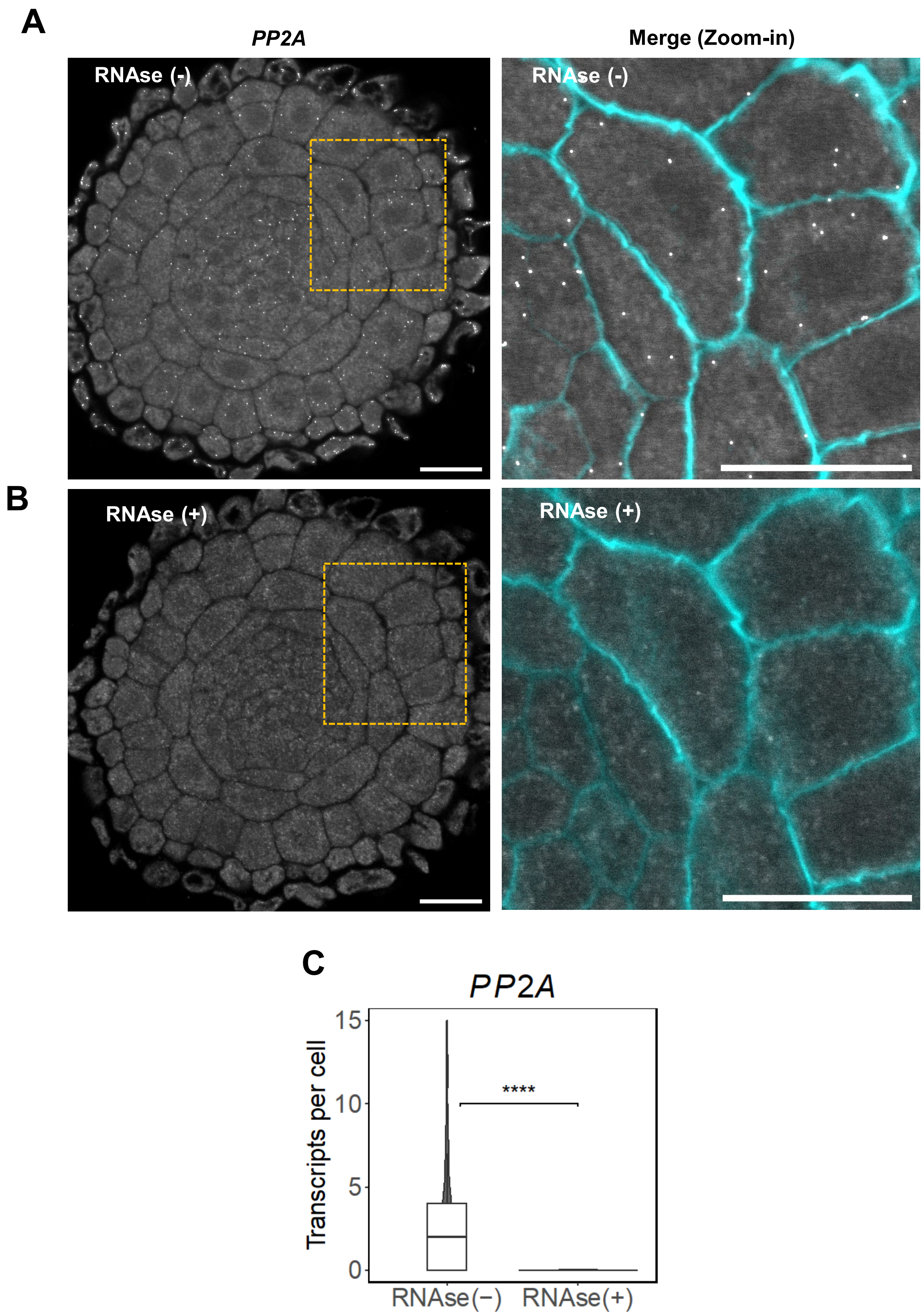


**Figure S2. *PP2A* mRNA detection and quantification using cryo-smFISH on *Arabidopsis* root cross-cryosections.** (A-B) Representative images of cryo-smFISH using probes targeting *PP2A* mRNA in *Arabidopsis* root cross-cryosections before (A) and after RNAse treatment (B)**.** Zoom-in images (right panels) of regions highlighted in yellow on the left panel. Discrete white dots correspond to individual RNA molecules. Cell walls stained with SR 2200 dye (cyan). Scale bars, 15µm.(C) Violin plot showing the number of transcripts per cell detected before RNAse (-) and after RNAse (+) treatment. Boxes inside indicate the interquartile range (IQR 25-75%), depicting the median values as a horizontal line. Whiskers represent the ±1.58xIQR value. A two-sided t-test was performed to compare both conditions. p-value less than 0.0001 was denoted by ****.

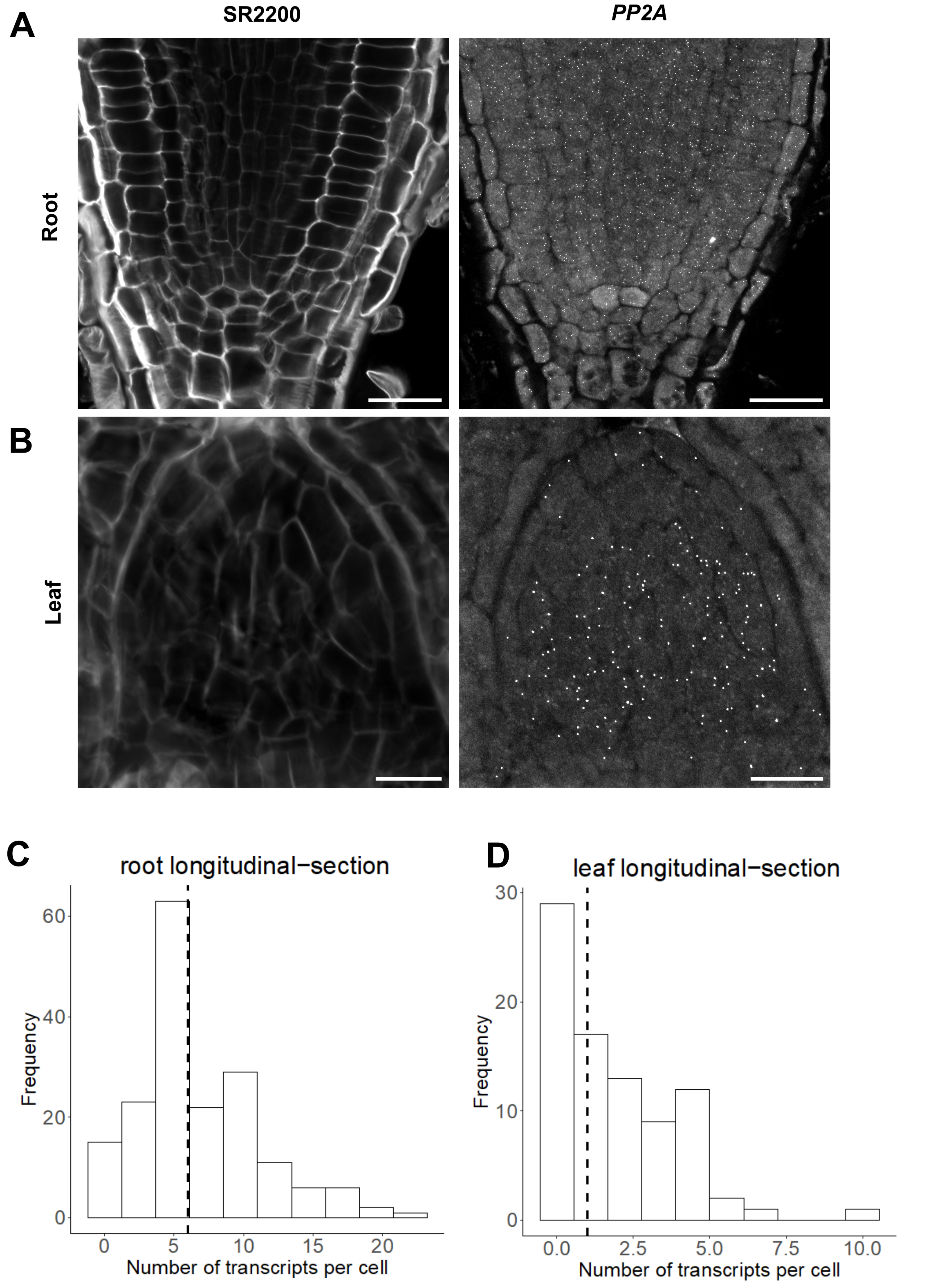


**Figure S3. *PP2A* mRNA detection in *Arabidopsis* longitudinal root and leaf cryosections.** (A, B) Representative confocal images of *PP2A* mRNA detected by cryo-smFISH on *Arabidopsis* root longitudinal-section (A) and young leaf longitudinal-section (B). Left panels: an overview of the tissue using celll wall SR 2200 dye; Right panel: *PP2A* cryo-smFISH signals (white dots) corresponding to individual mRNA molecules. Inset: inverted zoomed-in image from the from regions highlighted in yellow (box). Scale bars, 10µm. (C, D) Histograms showing the distribution of the number of transcripts per cell detected on root longitudinal-section (C) and leaf longitudinal-section (D). The dashed lines mark the median value.

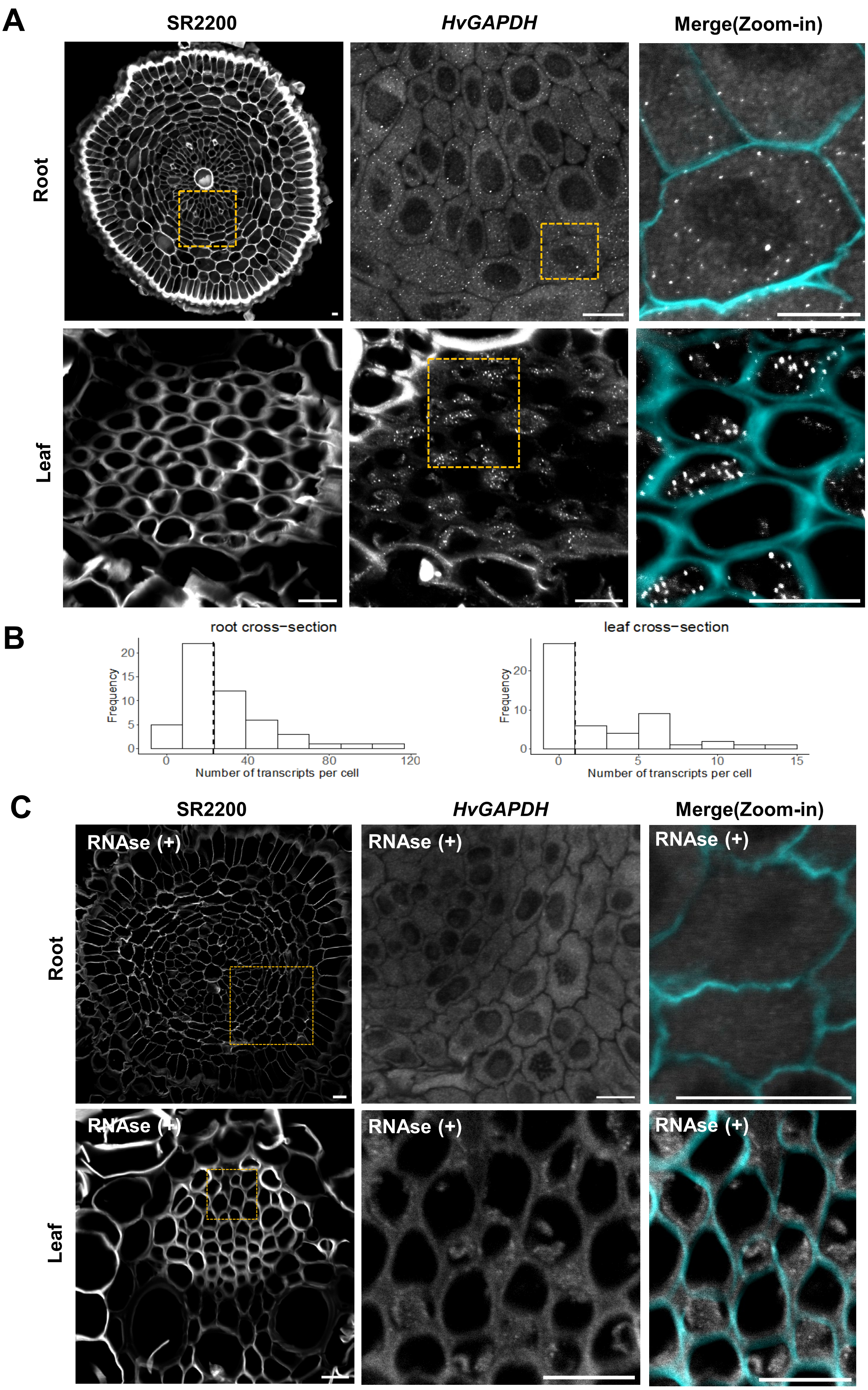


**Figure S4. Detecting *HvGAPDH* mRNA in barley roots and leaves using cryo-smFISH.**
(A, C) Representative images of cryo-smFISH using probes targeting *HvGAPDH* mRNA in barley root and leaf cross-sections with (C) and without (A) RNAse treatment**.** Left panels: cell wall staining with SR 2200 dye; Middle panels: cryo-smFISH channel; Right panels: zoomed-in images of regions highlighted in yellow in the left panels showing cryo-smFISH channel (white) and cell wall staining (cyan). Scale bars, 10µm. (B) Histograms showing the distribution of the number of transcripts per cell detected in the root (left) and leaf cross-sections (right). Dashed lines indicate the median value. Experiments were repeated independently 2 times.

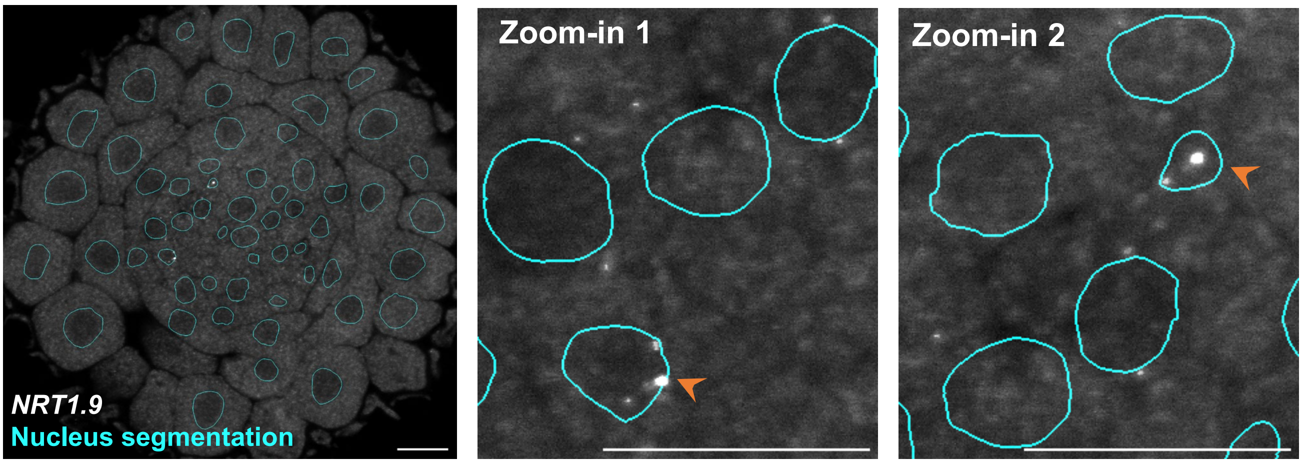


**Figure S5.** ***NRT1.9*** **expression pattern visualized by cryo-smFISH**
Representative images of cryo-smFISH using probes targeting *NRT1.9* mRNA in Arabidopsis root cross-cryosections. Middle and right panel: zoomed-in images showing *NRT1.9* transcripts (white dots). Nucleus outlines obtained from segmentation using DAPI (cyan). Scale bars, 10µm.

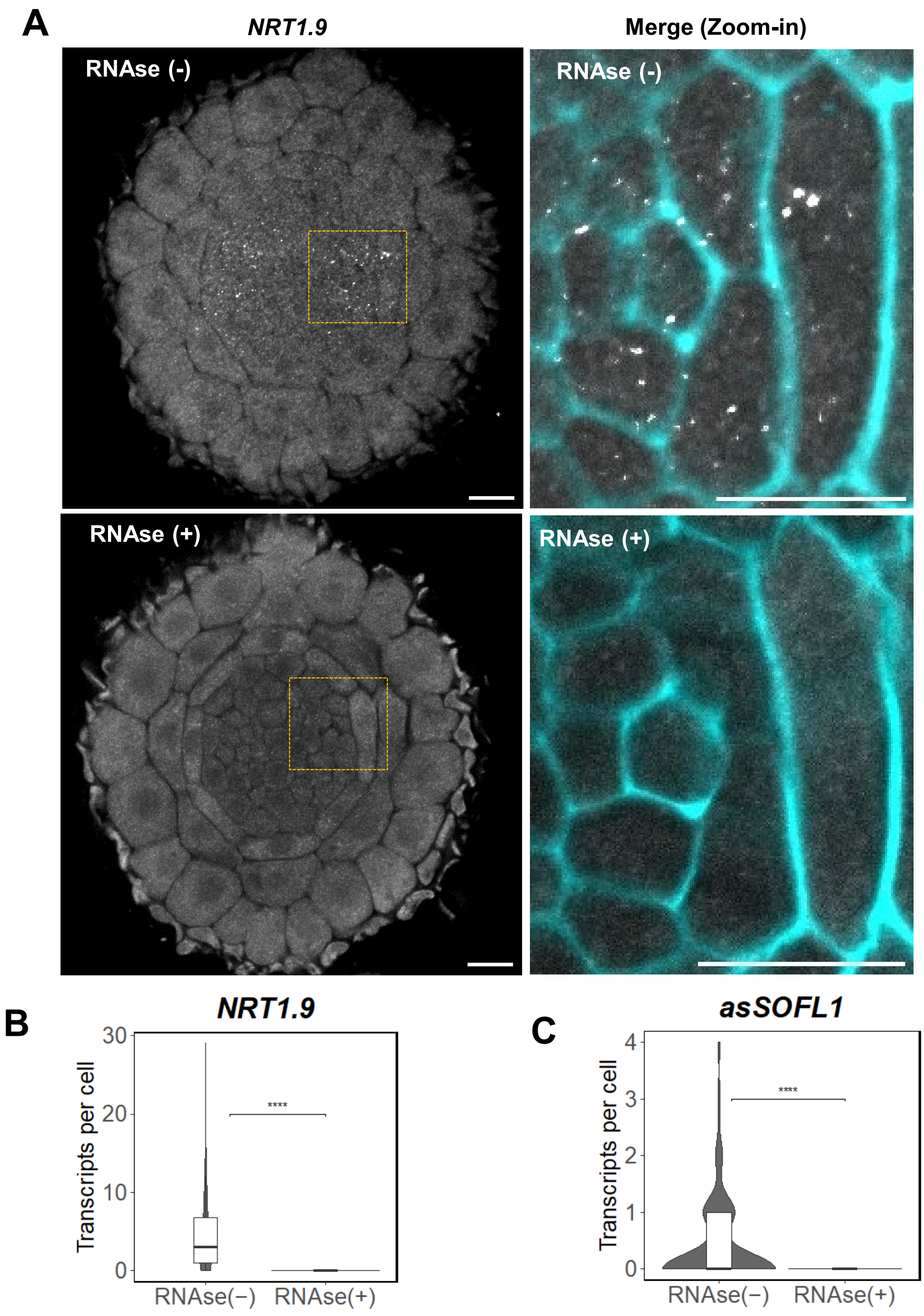


**Figure S6. RNAse A controls for cryo-smFISH experiments on *NRT1.9* and *asSOFL1*.**
(A) Representative images of cryo-smFISH using probes targeting *NRT1.9* mRNA in *Arabidopsis* root cross-cryosections before (-) and after (+) RNAse treatment**.** Zoom-in images (right panels) of regions highlighted in yellow on the left panel. Discrete white dots correspond to individual RNA molecules. Cell walls are stained with SR 2200 dye (cyan). Scale bars, 10µm. (B, C) Violin plots showing the distributions and quantification of the number of transcripts per cell before RNAse (-) and after RNAse (+) treatment for *NRT1.9* and *asSOFL1*. The dashed lines in histograms indicate the median value. Violin plots: boxes inside indicate the interquartile range (IQR 25-75%), depicting the median values as a horizontal line. Whiskers represent the ±1.58xIQR value. A two-sided t-test was performed to compare both conditions. p-value less than 0.0001 was denoted by ****.

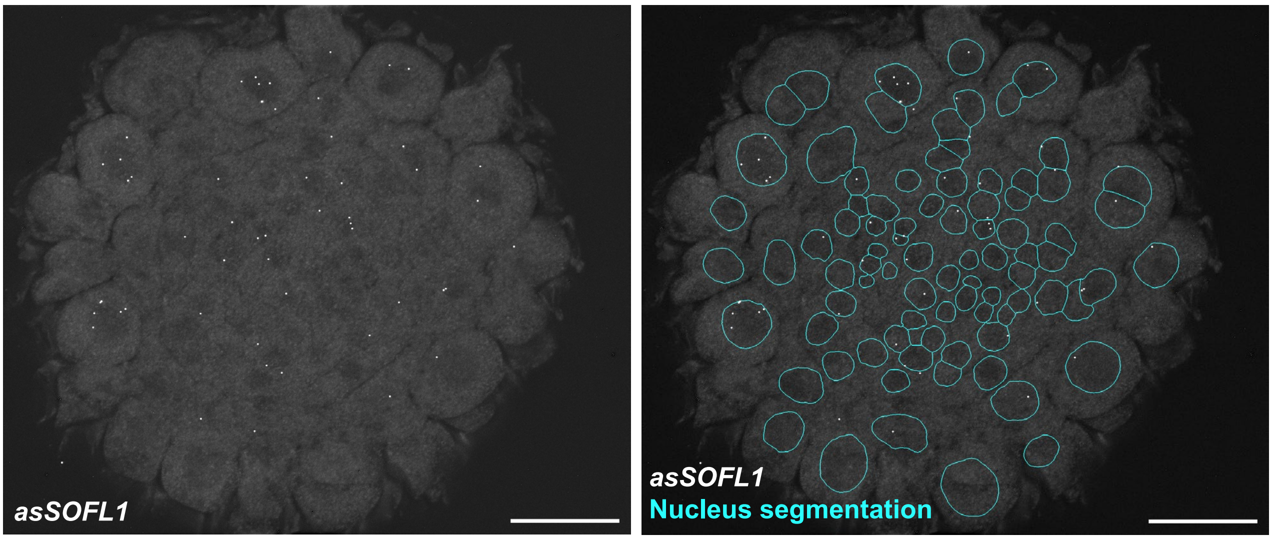


**Figure S7. *asSOFL1* expression pattern visualized by cryo-smFISH**
Representative Z project images of cryo-smFISH using probes targeting *asSOFL1* RNA in Arabidopsis root cross-cryosections. Middle and right panel: zoomed-in images showing *asSOFL1* transcripts (white dots). Nucleus outlines obtained from segmentation using DAPI (blue). Scale bars, 10µm.

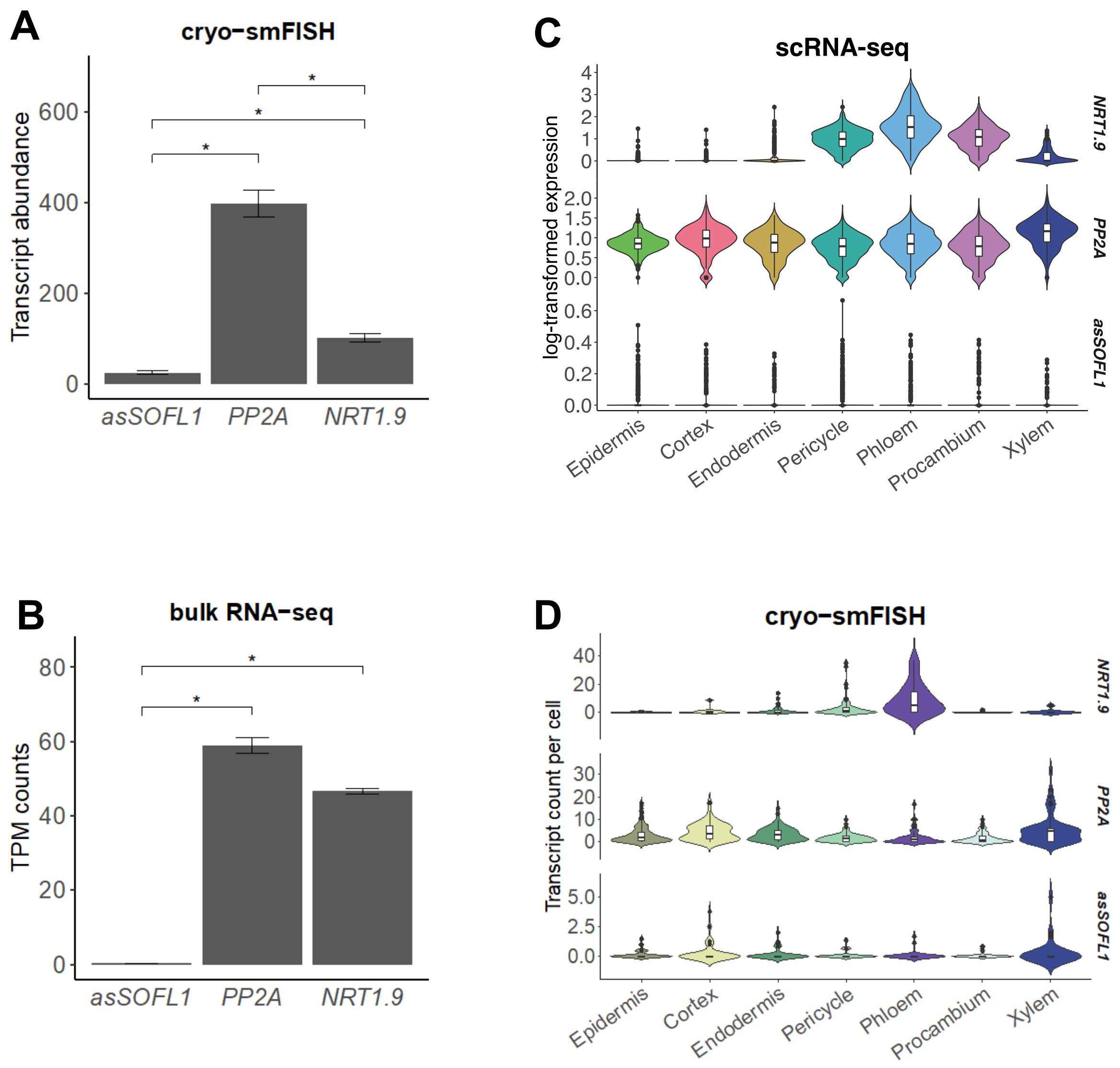


**Figure S8: Comparison of *NRT1.9*, *PP2A* and *asSOFL1* expression between scRNA-seq, smFISH and bulk RNA-seq.** (A, B) Bar plots showing the expression of *NRT1.9*, *PP2A*, and *asSOFL1* in the *Arabidopsis* root tip using cryo-smFISH (A) and bulk RNA-seq data (B). Error bars represent the mean ± SEM. Quantitative data for smFISH and bulk RNA-seq results originated from three replicates and two replicates, respectively. A two-sided t-test was performed to compare both conditions. A p-value greater than 0.05 indicates no statistical significance (not shown), while p-values less than 0.05, 0.001, and 0.0001 were denoted by*, **, and ***, respectively.(C, D) Violin plots showing the expression of *NRT1.9*, *PP2A,* and *asSOFL1* in individual cells detected by scRNA-seq (C) and cryo-smFISH (D). The number of transcripts is normalized by the number of cells within the cell type. The plotting area is scaled by width. Violin plots: boxes inside correspond to the interquartile range (IQR 25-75%), and horizontal lines indicate the median values. Whiskers show the ±1.58xIQR value. Statistical analyses were performed with one-way ANOVA followed by Tukey's honestly significant difference (HSD) tests.

**Table S1. Cryo-smFISH strenghts and limitations.**

| | |
|---|---|
| **Strengths** | - Detection and visualization of single molecules of RNAs with cellular and sub-cellular resolution in plant tissue sections;<br>- Easy to implement and relatively fast compared to other sectioning methods;<br>- Possibility of performing smFISH on very thick specimens;<br>- Images can be acquired with widefield and confocal microscopes;<br>- Can be combined with immunofluorescence for co-detection of RNA and protein in the same cells (antibody penetration is more easily achieved than with whole-mounted preparations);<br>- Can be used to validate single-cell RNA seq. No transgenic lines are needed;<br>- Can be implemented in non-model species. |
| **Limitations** | - Preservation of tissue structure can be challenging for certain tissues (ex. *Arabidopsis* roots) and requires optimization and practice;<br>- Cannot be used to study more than 4 genes at the same time due to fluorophore limitations (i.e., only a small number of colors can be used for microscopy);<br>- Cannot be used to study RNA that are shorted than ~600nt in length (probes are 18-22 mers DNA oligonucleotides and a minimum of 25-30 probes is required per target). |

**Table S2. smFISH probe sequences used in this study and the corresponding dyes (Quasar570 or Qusar670).** Probe Sequences (5'-3').

| Probe nr | *PP2A* (Quasar570) | *HvGAPDH* (Quasar570) | *NRT1.9* (Quasar670) | *asSOFL1* (Quasar670) |
|---|---|---|---|---|
| 1 | ccgagcgatctatcaatcag | ggttttataaagccaggagc | caaatgtctcatttcctacc | gaccagctgaattcagtct |
| 2 | gacatcctcaccaaaactca | agaggtggggagggtttaag | cagattcgatgaacttccaa | cgaatccagaggctttctt |
| 3 | tcgggtataaaggctcatca | agtgtctggaagcggatatg | cagccgtgatgctcttcat | gcaacgaaagccaccaagt |
| 4 | tagctcgtcgataagcacag | cgaaggagtggaactgacgc | ccgtagatgttgacgacttt | atcacaagcgacgtttcca |
| 5 | ccaagagcacgagcaatgat | cgatcttaatcttgcccatg | gatggtgccaaagttgcttg | aggtccacattaaacccac |
| 6 | atcaactcttttcttgtcct | atccttccgaaaccgttgat | gagtcacagaggaaagcagc | gtttctcggaggcacaatc |
| 7 | catcgtcattgttctcacta | tgatgaaggggtcgttgacg | aagatagggtcttgtagcgg | tccttttgtctgattccat |
| 8 | atagccaaaagcacctcatc | catgtacgtcatgtactcgg | agaaagcaagcgatcatggc | atccacagccacacacaaa |
| 9 | atacagaataaaaccccca | cgtgaactgtgtcgtacttg | cgaaccctttgcatcaaaca | cgagtttcttcgtaccgat |
| 10 | caagtttcctcaacagtgga | atgtcactgtgcttccaatg | gatgcaggggatgaatcaca | tgaagctactacattgccc |
| 11 | tcatctgagcaccaattcta | agcgtcttgtcgtctttgag | tatttccttagcgcattgag | gttcactccatttctggat |
| 12 | tagccagaggagtgaaatgc | cctgacgccaaagacagtaa | tggcccattgcacacgctt | tgcttgtttgcatcactca |
| 13 | cattcaccagctgaaagtcg | gtggactccacaacgtaatc | gaaacatgatttgtccgatg | atcctctgctgatcacatg |
| 14 | ggaaaatcccacatgctgat | ttgtccttgtcagtgaagac | accagcaaaaccattgctcc | accattcctgaattgcagg |
| 15 | atattgatcttagctccgtc | cagagatgaccaccttcttg | atggcctaatcccaccggct | gagtcttggactagccata |
| 16 | attggcatgtcatcttgaca | aaacataggggcatctttgc | gatcagcaccaaatggaag | agatgtgcacgttcggtag |
| 17 | aaattagttgctgcagctct | ttgtcctcattgacaccaac | ttcttttgtcttcggatcga | ttcctgctagttacaagct |
| 18 | gctgattcaattgtagcagc | atgttaacatccgaggtgta | aaactctcaatccctcgttt | aatagcatgtgcacgttcc |
| 19 | ccgaatcttgatcatcttgc | gtgcagctagcatttgagac | gtgaaggtgaagaaatacc | cccaaccttcacgaagatt |
| 20 | caaccctcaacagccaataa | taatgaccttagctaggga | cgatgagggttaacgacacc | tgttattctcaggtcgctt |
| 21 | ctccaacaatttcccaagag | tcagaccctcaataatacca | cagctcacgtttgactgaac | aagaggaagaccctcacgt |
| 22 | caaccatataacgcacacgc | tgatggcatgaacagtggtc | agctgggatggctaaacca | acatagctggcaactttcc |
| 23 | agtagacgagcatatgcagg | catcaacggtcttctgtgtg | tgcaaccaagtagcattag | gcatatcctgataagcgca |
| 24 | gaacttctgcctcattatca | gctggggatgatgttaaagc | gagcttagaaccagcgaaga | tcttcagagacttccttct |
| 25 | cacagggaagaatgtgctgg | cattcagctcaggaagaacc | ccactagctttaaccttgac | agttatctgcacggttctc |
| 26 | tgacgtgctgagaagagtct | ccggaaagacataccggtaa | acacgagttatgctgtgaa | ttttgtagaagctgtccca |
| 27 | cccattataactgatgccaa | aactgacacatccacagtgg | cgcctcttcttgatcgcgac | attgttttcttttgccgga |
| 28 | tggttcacttggtcaagttt | cggttctaacagtgagatca | aagctcgttaggcccaacgg | agccatgtgtaggtagttg |
| 29 | tctacaatggctggcagtaa | catcatatgatgcagccttc | tggcccaatttcgagttct | caatcgtgttgtttgcctt |
| 30 | cgattatagccagacgtact | ccttgatagccttcttgatg | ttagtacctgaactgctctg | gcaatcagcttccaaagca |
| 31 | gactggccaacaagggaata | catgatacccttgagctttc | actaagagaagtagtgagg | gtgatgattccatggcttc |
| 32 | catcaaagaagcctacacct | accaaatcttcctcaacgta | ttggattgccgatttgtcaa | agacgccttccatggaaat |
| 33 | ttgcatgcaaagagcaccaa | tgtcaccaacgaagtcggtg | catctttgttcagcttgtcg | acagcagttgtgagtcagg |
| 34 | acggattgagtgaaccttgt | agccttagcatcaaagatgc | gtttccatgcatccaccggc | gaggctctgaagaggagga |
| 35 | cttcagattgtttgcagcag | cgaaatggtcgttcagagca | ctccacttgttgcatactg | ccatggagtctccgagaaa |
| 36 | ggaccaaactcttcagcaag | tgtcataccacgagacaagc | acccggatcacgcatttaac | cttggttttgttttcctct |
| 37 | ggaactatatgctgcattgc | cgacaacacggttgctgtaa | taaggcggccgagagccaaa | tatgaggcttcctataggc |
| 38 | gtgggttgttaatcatctct | ggaaagcagaacgctctact | gttgaatgtaagctaggtag | ggttgccatgtggtgaaac |
| 39 | tgcacgaagaatcgtcatcc | catagacaaaggggcacctc | ctgaaagattgtgtaggttg | tttgtctggtagtgcttca |
| 40 | ttactggagcgagaagcgat | gacaccatccacatttattc | aggcgtctatcggattgaag | gaagggaggcactaacctt |
| 41 | gaacatgtgatctcggatcc | aacatgaaccaggcgtctag | ggatctggaaacttcctgga | tgtttgttcatgtgggtct |
| 42 | ctctgtctttagatgcagtt | cagatggtaactcatgtcca | aagacggtataagaaccggc | ggcctccactgaaagaaca |
| 43 | catcattttggccacgttaa | acttgactagcaactcggtg | tgaatatcgtcattccaagc | tgatttgcaagctgtcacc |
| 44 | cgtatcatgttctccacaac | cacttactcaggcaaacaga | tacaaggacgcggtcataga | caaattctacacgcccact |
| 45 | atcaacatctgggtcttcac | caaacacttactcaggctgt | gcctgtatactttctaagga | gagtccaactccattcttt |
| 46 | ttggagagcttgatttgcga | ggatcgcagcatcaaacatc | tctctgcagttgtgtgatac | gtgacattccacagtcaca |
| 47 | acacaattcgttgctgtctt | gacgggagtagtactcaaca | atgcatagaaacagtcctgc | cccgatgatgctttcgaat |
| 48 | cgcccaacgaacaaatcaca | cgctccgtcctataatataa | tgcagacaccatcatactt | caagttccatccttagtct |
